# PALM-Seq: integrated sequencing of cell-free long RNA and small RNA

**DOI:** 10.1101/686055

**Authors:** Xi Yang, Taifu Wang, Sujun Zhu, Juan Zeng, Yanru Xing, Qing Zhou, Zhongzhen Liu, Haixiao Chen, Jinghua Sun, Liqiang Li, Jinjin Xu, Chunyu Geng, Xun Xu, Jian Wang, Huanming Yang, Shida Zhu, Fang Chen, Wen-Jing Wang

**Affiliations:** BGI-Shenzhen, Shenzhen 518083, China; Obstetrics Department, Shenzhen Maternity and Child Healthcare Hospital, Shenzhen, Guangdong Province, China; China National GeneBank, BGI-Shenzhen, Shenzhen 518120, China; BGI Education Center, University of Chinese Academy of Sciences, Shenzhen 518083, China; MGI, BGI-Shenzhen, Shenzhen 518083, China

**Author notes:** These authors contributed equally: Xi Yang, Taifu Wang, Sujun Zhu.

## Abstract

Cell-free RNA, including both long RNA and small RNA, has been considered important for its biological functions and potential clinical usage, but the major challenge is to effectively sequence them at the same time. Here we present PolyAdenylation Ligation Mediated-Seq (PALM-Seq), an integrated sequencing method for cell-free long and small RNA. Through terminal modification and addition of 3’ polyadenylation and 5’ adaptor, we could get mRNA, long non-coding RNA, microRNA, tRNA, piRNA and other RNAs in a single library. With target RNA depletion, all these RNAs could be sequenced with relatively low depth. Using PALM-Seq, we identified pregnant-related mRNAs, long non-coding RNAs and microRNAs in female plasma. We also applied PALM-Seq to sequence RNA from amniotic fluids, leukocytes and placentas, and could find RNA signatures associated with specific sample type. PALM-Seq provides an integrated, cost-effective and simple method to characterize the landscape of cell-free RNA, and can stimulate further progress in cell-free RNA study and usage.

## Introduction

After cell-free RNA (cfRNA) was discovered, its importance has been proven. CfRNA plays important roles in cell communications^1,2^, and its biogenesis, uptake^3^ and distribution^4^ are correlated with physical and pathological situations^5^. Besides this, cfRNA carries information from human tissues^6,7^ or tumor^8,9^. According to these reasons, cfRNA is considered as potential targets for disease interventions and useful biomarkers for disorder prediction.

CfRNA in plasma is usually made up of degradative small fragments with size smaller than 200nt, very low concentration (may lower than 10 ng/mL)^10^, and different terminal modification^11,12^, and these properties make it difficult to research. Since 2014, next-generation sequencing was used for cfRNA studies^13,14^, but some problems were not solved till now, including requirement of large volume for blood^15^, no uniform sequencing method for all cfRNA fractions^12^, high cost for large scale cfRNA library preparation, and as well as low mapping rate.

Here, we presented a new method, PolyAdenylation Ligation Mediated-Seq (PALM-Seq), to get all sorts of RNA in one library with lower cfRNA input. Additionally, PALM-Seq provided a new possible way for entire transcriptome sequencing with low input of RNA samples. This approach could enable us to get more comprehensive results for large population studies and clinical usages of cfRNA.

## Results

### Principle of PALM-Seq

For amount of cfRNA was low, in order to avoid RNA degradation or adsorption to the tube, 3’ polyadenylation was added by E. coli Poly(A) Polymerase (PAP)^16^ in the first step. The terminal modification of cfRNA was different, however T4 Polynucleotide Kinase (T4 PNK) could make most part of them to 5’ end phosphorylated and 3’ end hydroxyl^17,18^ except those with 5’ cap. Because T4 PNK and PAP could work in the same buffer with ATP, separate step of pretreatment for cfRNA with T4 PNK was unnecessary. Then, 5’ adaptor was then ligated by T4 RNA Ligase 1^19^. In contrast to 3’ DNA adaptor used in common small RNA seq, 3’ polyadenylation could not be degraded by DNase I which was necessary for RNase H method^20^. So, rRNA or other uninterested, abundant RNA could be easily removed by RNase H method. DNase I treatment could also prevent the possible contamination of cell-free DNA. Finally, RNA with 3’ polyadenylation and 5’ adaptor was reverse transcribed by oligo(dT) with 3’ adaptor and amplified by PCR (Fig. 1a). Most part of the library was short fragments as expected. The processing pipeline of sequencing data was shown in Fig. 1b.

**Figure 1.**
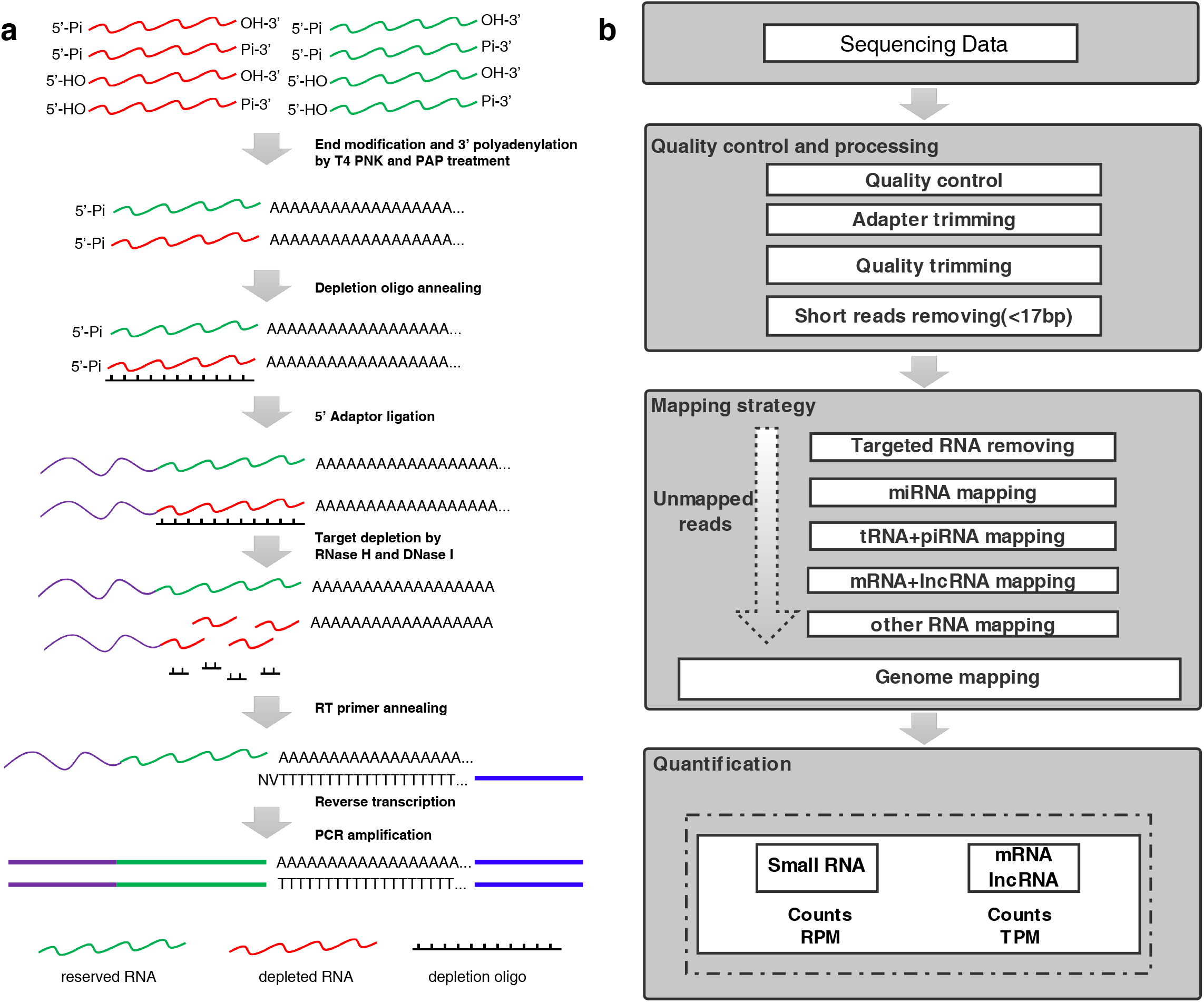
Scheme of PALM-Seq. **a**, Principle of PALM-Seq. Cell-free RNAs (cfRNA) are terminal modified and added with 3’ poly(rA) tail and 5’ adaptor. Targeted RNAs are depleted. Finally, RNAs are converted to cDNA by reverse transcription and PCR amplification. **b**, Data Analysis. Low quality data are removed at first, and then adapter trimming is made. Reads shorter than 17bp, rRNA and/or other targeted RNA (e.g. Y RNA and Vault RNA in plasma) reads are filtered. The mapping is performed in the order of miRNA, tRNA/piRNA, mRNA/lncRNA and other RNA. The unmapped reads will be mapped to genome if necessary. Small RNAs are quantified by counts and Reads Per Million (RPM), and mRNA or lncRNA are quantified by counts and Transcripts Per Million (TPM).

In order to test the effect of T4 PNK treatment (PNKT) and target depletion (TD) which was performed through RNase H method, and identify the best work condition for cfRNA, we used four samples, each of which was treated in four work conditions, with or without PNKT and with or without TD. We also wanted to know whether TD could remove other abundant RNA, so Y RNA and Vault RNA which were rich in plasma cfRNA^21^ and not interested in this study, were also depleted for plasma samples.

### PALM-Seq could capture plasma mRNA, lncRNA and small RNA with high complexity

Complexity is especially important for low-quantity and low-quality RNA sequencing^20,22^, and it is better to get more kinds of RNA in one library. To assess the performance of different treatments and methods, we analyzed the data to account for differences in number of different genes detected and contribution of different kinds of RNA. PALM-Seq with PNKT (TD or no TD) could detect larger number of mRNA (Fig. 2a) or lncRNA (Fig. 2b), and it could also detect miRNA (Fig. 2c), tRNA (Fig. 2d) and piRNA (Fig. 2e), while the fewer miRNAs were covered. PALM-Seq with no PNKT and no TD were more efficient to detect miRNA (Fig. 2c), tRNA (Fig. 2d) and piRNA (Fig. 2e), but it covered less mRNA (Fig. 2a) and LncRNA (Fig. 2b).

**Figure 2.**
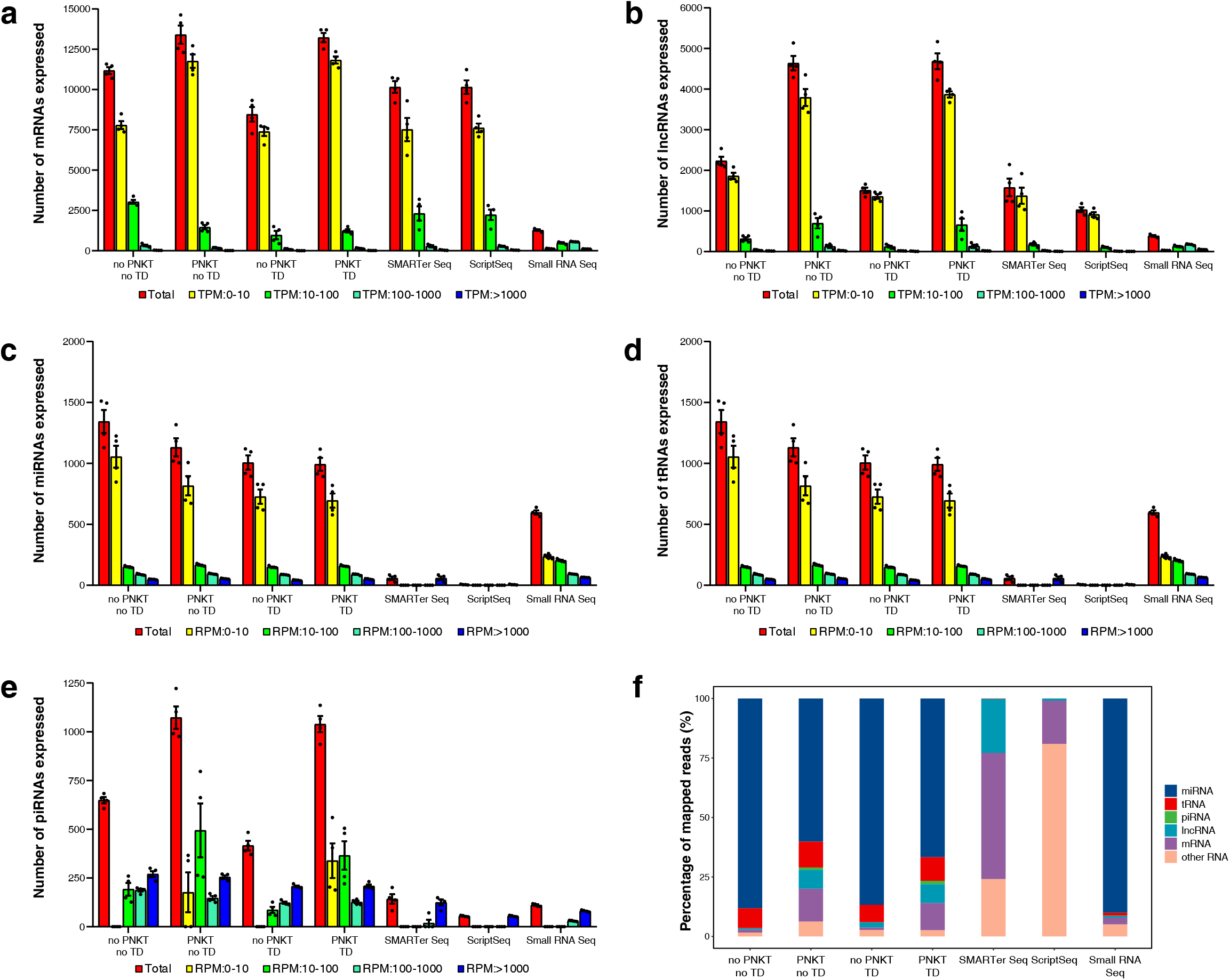
Complexity and RNA biotype contribution of different RNA-seq methods for plasma cfRNA. **a-e**, Number of mRNAs (**a**), lncRNA (**b**), miRNA (**c**), tRNA (**d**) and piRNA (**e**) expressed in each treatment or method. Four samples are used for each subgroup, the results are shown as mean ± S.E.M. (Standard Error of Mean). **f**, Average ratio (n=4) of mapped reads for each type of RNA in different cfRNA sequencing methods. PNKT: T4 polynucleotide kinase treatment. TD: Target depletion.

To further qualify our method, we referred to public data produced by SMARTer Seq (by using Clontech SMARTer Stranded Total RNA-Seq Kit - Pico)^15^, ScriptSeq (by using Illumina ScriptSeq v2 Kit)^23^ and Small RNA Seq (by using NEBNext Small RNA Library Preparation Kit)^24^ to quantify if PALM-seq was comparable or even better. While SMARTer Seq and ScriptSeq were designed for large fragment RNA, they were effective to detect mRNA or lncRNA (Fig. 2a-b), but they could not cover small RNA (Fig. 2c-e). In contrast, Small RNA Seq performed well in detecting miRNA (Fig. 2c), but was not suitable for lncRNA or mRNA (Fig. 2a-b). RNA ratio also showed that mRNA and lncRNA were rich in PALM-Seq with PNKT (TD or no TD), SMARTer Seq or ScriptSeq, however miRNA was rich in PALM-Seq with no PNKT (TD or no TD) or Small RNA Seq (Fig. 2f). In addition, tRNA and piRNA ratio was higher in PALM-Seq (Fig. 2f).

Length contribution of different sort of RNA were also checked to further estimate the effect of PNKT. Large fragments of mRNA or lncRNA could be captured without PNKT, however, more small fragments of mRNA or lncRNA would be captured with PNKT, for small fragments might be produced by RNase (e.g. RNase A) with 3’ phosphorylated and 5’ hydroxyl. Length contribution of small RNA were less influenced by PNKT. The information provided by small fragments of mRNA or lncRNA would be important especially for small amount or long-stored samples without enough long fragments, so PNKT was acquired for mRNA or lncRNA analysis.

These results illustrated that PALM-Seq could be used for miRNA, tRNA and piRNA whether with or without PNKT, but PNKT was necessary for mRNA and lncRNA analysis.

### TD could remove uninterested RNA efficiently and improve mapping rate

Mapping rate is important for data analysis to detect more genes, splicing or other variations with lower sequencing depth^20^. Depletion of uninformative abundant RNA (e.g. rRNA) can increase mapping rate and maximize the coverage of other transcripts present in a sample. To test the efficiency and necessity of TD, we calculated rRNA ratio (Fig. 3a). We found that rRNA could be removed efficiently by TD, especially in PNKT condition, otherwise, the rRNA ratio would be extremely high. In no PNKT condition, TD was dispensable. In addition, Y RNA and Vault RNA could also be removed by TD. SMARTer Seq removed rRNA through CRISPR, so the rRNA ratio was low, but ScriptSeq or Small RNA Seq data contained high ratio of rRNA.

**Figure 3.**
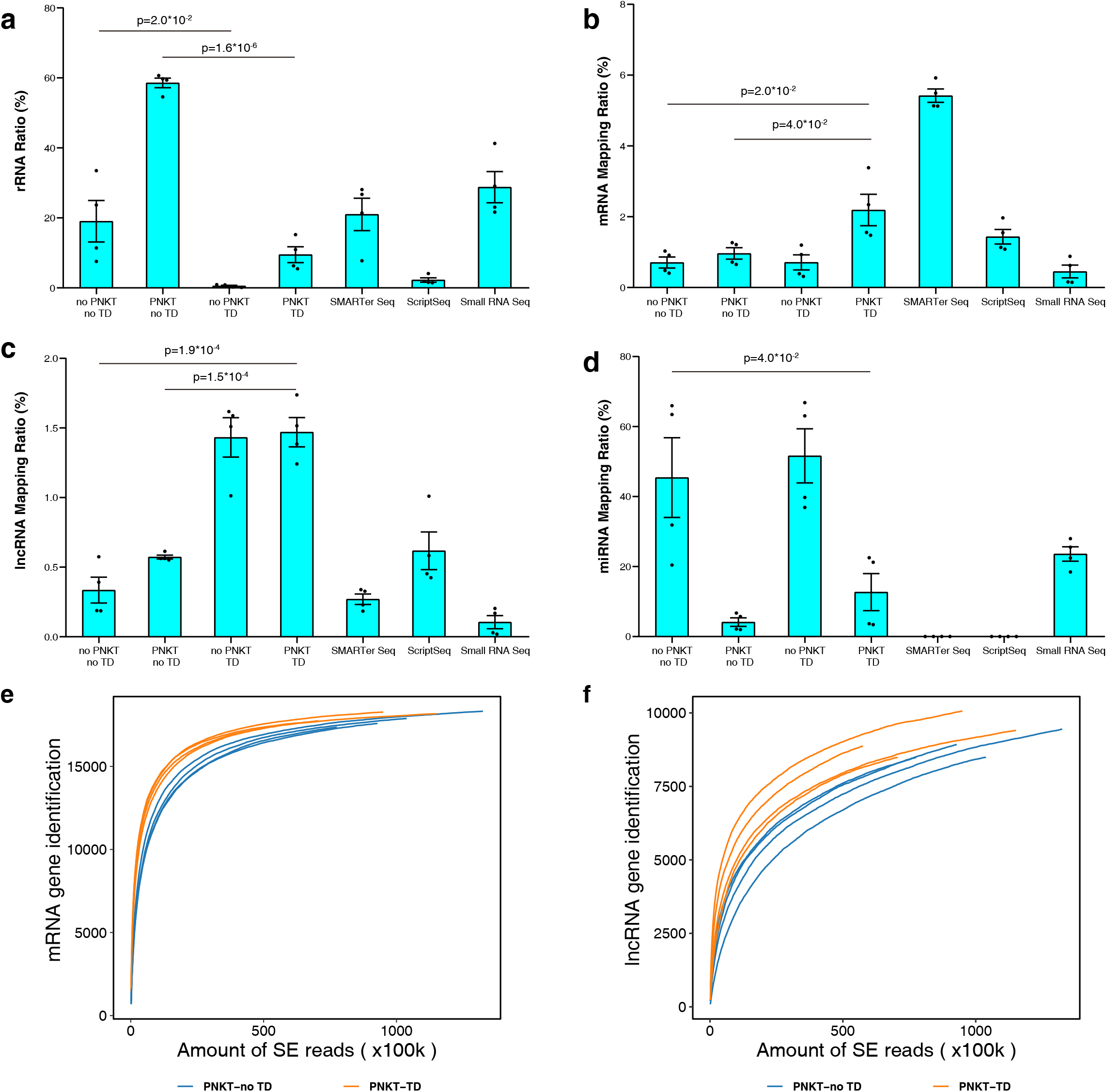
Mapping rate of different cfRNA sequencing methods. **a-d**, Performance of each library with respect to rRNA ratio (**a**, normalized to read number after low-quality reads and <17bp reads have been filtered), mRNA mapping rate (**b**, normalized to total read number), lncRNA mapping rate (**c**, normalized to total read number) and miRNA mapping rate (**d**, normalized to total read number). Unpaired two-sided Student’s t-test is used to assess significance. The original p values are shown here. **e-f**, Saturation plots for mRNA (**e**) and lncRNA (**f**) shows the influence of TD to the acquired sequencing depth when PNKT is used.

To further estimate informative data ratio, we calculated mRNA, lncRNA and miRNA mapping rate to total reads (Fig. 3b-d). We found that TD increased mRNA and lncRNA mapping ratio (Fig. 3b-c) and decreased the acquired sequencing depth^25^ when PNKT was used (Fig. 3e-f). However, TD did not significantly influence miRNA mapping ratio.

Then we compared mapping rate among different methods. For mRNA and lncRNA mapping rate, PALM-Seq with PNKT and TD performed similarly with SMARTer Seq, and much better than Small RNA Seq. ScriptSeq performed best for mRNA but not well for lncRNA. For miRNA mapping rate, PALM-Seq with PNKT and TD performed similarly with Small RNA Seq, although lower than PALM-Seq with no PNKT and no TD.

These results suggested that TD was helpful, and it was necessary when PALM-Seq was used to detect both mRNA and miRNA, for rRNA ratio might be unacceptable if only PNKT was used.

We recommended PALM-Seq with PNKT and TD for the studies which needed both mRNA/lncRNA and miRNA, and PALM-seq with no PNKT and no TD for the researches focused on miRNA.

Because both mRNA/lncRNA and miRNA were needed in our further work, PALM-Seq with PNKT and TD were used for other samples mentioned in this study.

### PALM-Seq was a robust method for plasma RNA-Seq

An ideal cfRNA sequencing method should not only display high yield, complexity and mapping rate, but also have high reproducibility^22^. We tested the reproducibility of PALM-Seq and found that PALM-Seq with PNKT and TD showed the similar consistency for mRNA as SMARTer Seq or ScriptSeq (Fig. 4a), but the consistency of lncRNA was higher for PALM-Seq (Fig. 4b). PALM-Seq with PNKT and TD or PALM-Seq with no PNKT and no TD gave more stable results compared with Small RNA Seq (Fig. 4c).

**Figure 4.**
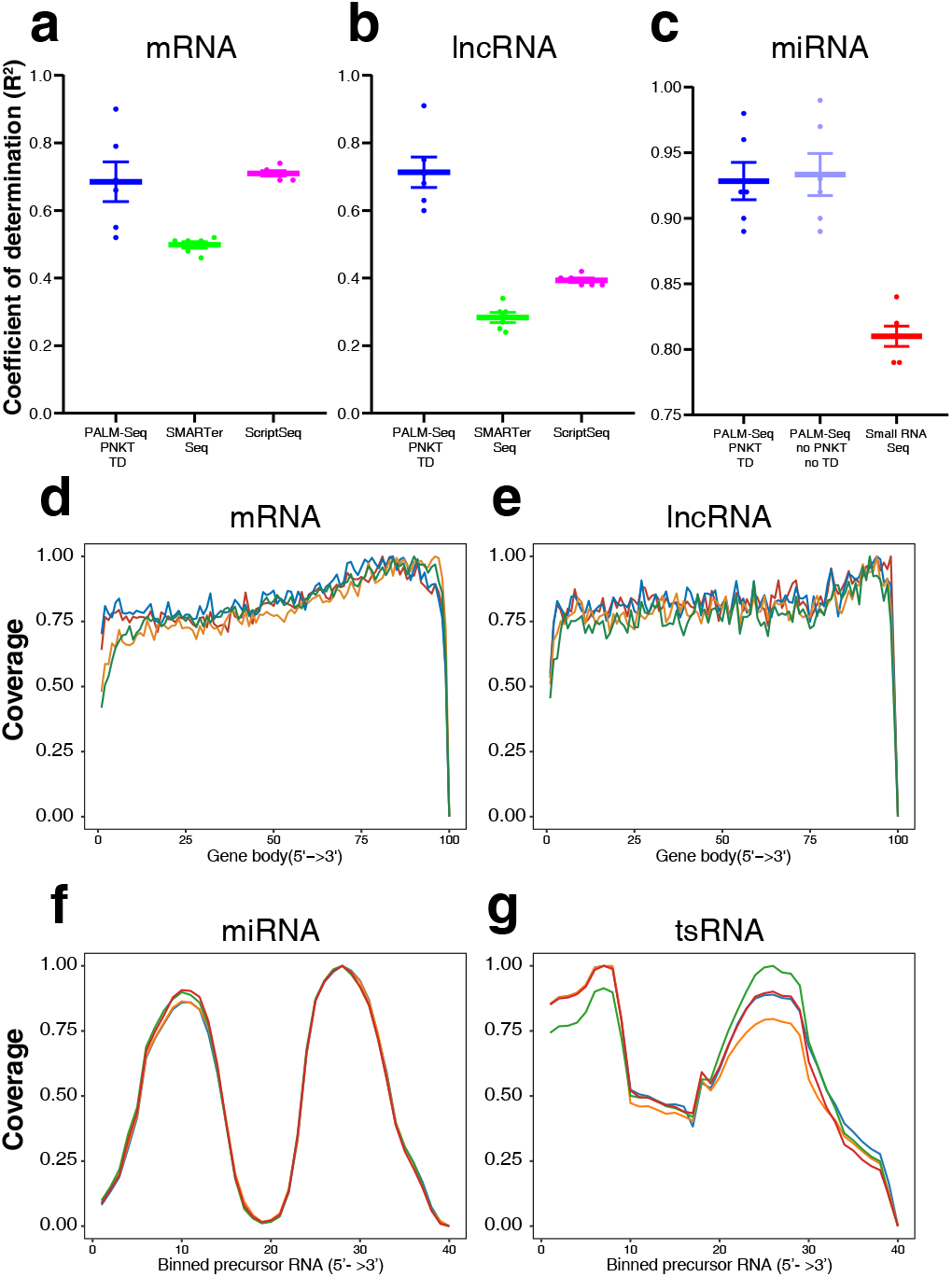
Reproducibility and transcript coverage of PALM-Seq. **a-c**, Reproducibility is measured by coefficient of determination (R^2^). PALM-Seq with PNKT and TD is compared with SMARTer-Seq or ScriptSeq for mRNA (**a**) and lncRNA (**b**). PALM-Seq with PNKT and TD or with no PNKT and no TD are compared with Small RNA Seq for miRNA (**c**). R^2^ is calculated through simple linear regression of log_2_(TPM+1) for mRNA and lncRNA or log_2_(RPM+1) for miRNA. d-e, Normalized coverage by position for mRNA (**d**) and lncRNA (**e**). For each library, the average relative coverage is shown at each relative position along the transcripts’ length. **f-g**, Coverage of miRNA (**f**) and tRNA-derived small RNAs (tsRNA, **g**) across the length of the precursor RNAs (divided into 40 bins). Different curves stand for different samples using PALM-Seq with PNKT and TD.

Coverage metrics might be critical for certain measurements. We calculated the variation in coverage of mRNA and lncRNA from 5’ to 3’ ^20^, and PALM-Seq did not show significant bias from 5’ to 3’ by considering all detected genes. We then aligned the small RNAs to their respective precursors^26^. As expected, the mature miRNAs were aligned to the two arms of the pre-miRNA hairpins (Fig. 4f). The tRNA-derived small RNAs (tsRNAs) were mainly derived from the two regions of the mature tRNA, but it showed slight difference with cell tsRNAs^26^ (Fig. 4g).

To estimate the failure rate of this method, we totally built 120 libraries for plasma cfRNA, and 2 of them failed for low yield (<300ng).

For further usage of PALM-Seq, we assessed the influence of sequencing depth^25^ and read length. First, we evaluated saturation plots for different transcript biotypes. It was shown that 20M clean reads (about 40M total reads) were enough for mRNA, lncRNA, miRNA and tRNA quantification. Due to the low piRNA ratio, more reads might be acquired for piRNA analysis. We then compared expression results with single read 35 cycles or 100 cycles, and there was no significant difference, because most of cfRNA fragments were short.

### Placental-specific genes could be detected in plasma cfRNA from pregnant women

One of the basic usages for RNA-Seq is differentially expressed gene (DEG) analysis^27^, so PALM-Seq was tested for DEGs. During pregnancy, placenta releases nucleic acids into blood. To test whether PALM-Seq could capture placental specific RNA, we collected blood samples from 4 nonpregnant females and 4 pregnant females (three time points: 12, 24 and 36 gestational week).

Principal components analysis^28^ (PCA) showed that there were no significant outliers. Then, we checked the expression of placenta-specific mRNA/lncRNA (Fig. 5a) or miRNA (Fig. 5b). As expected, these genes were only expressed in plasma from pregnant women. We verified 5 mRNA/lncRNA genes and 5 miRNAs. Protein coding gene such as *CGA, CSH1* and *CSH2*, and long non-coding gene such as *PLAC4*^15^, were very low or undetected in nonpregnant females, and increased during pregnancy. *S100A8* which are related to pregnant immunomodulation^15^, also increased (Fig. 5c). Placental-specific C17MC miRNA^29^ such as mir-526b-5p, mir-519d-3p, mir-515-5p and mir-512-3p, also existed in plasma of pregnant females; and we found mir-454-3p decreased (Fig. 5d). These results were validated by RT-qPCR (Fig. 5c-d). Together with the results above, it was suggested that PALM-Seq could provide reliable results for DEG analysis.

**Figure 5.**
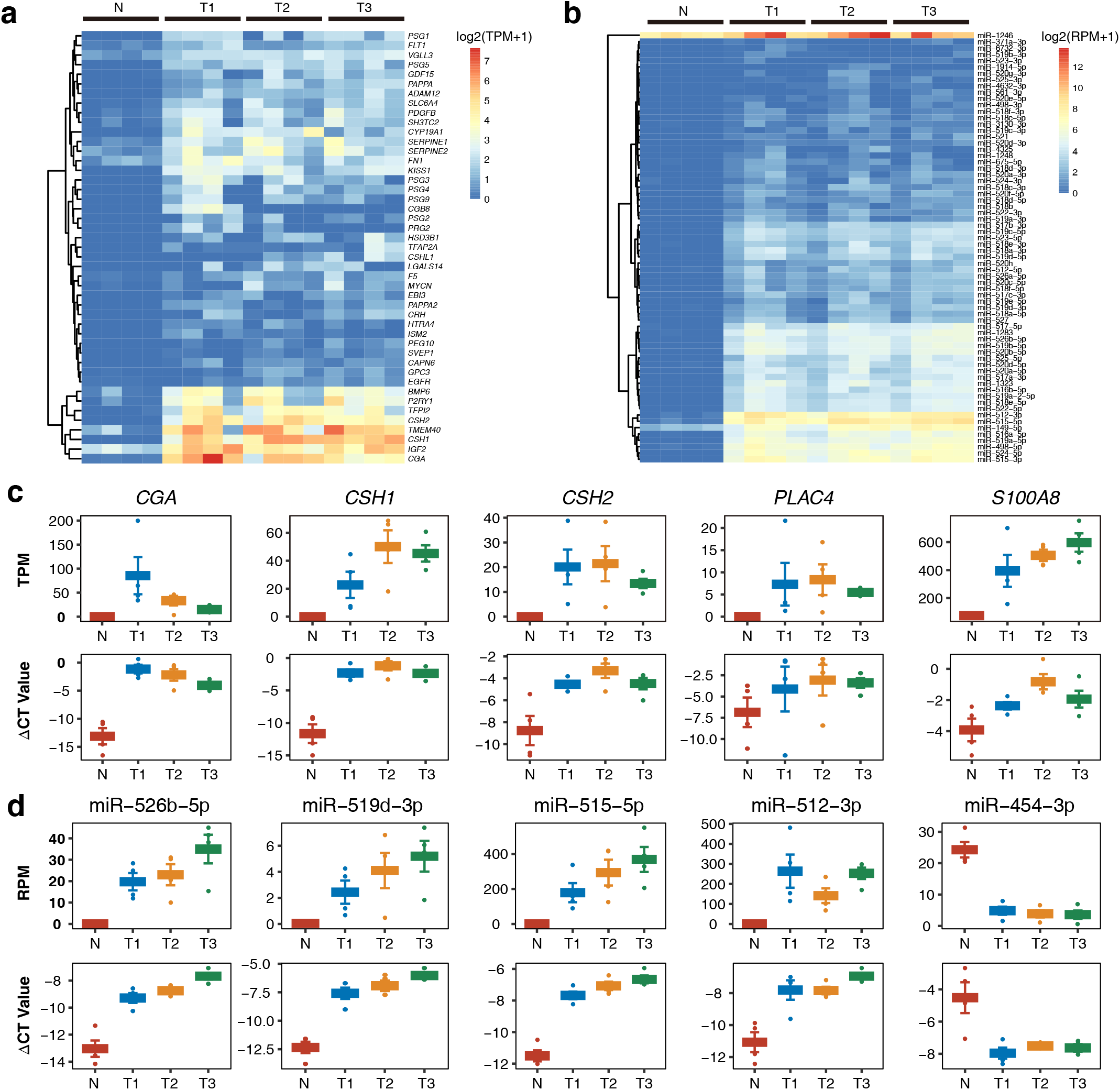
PALM-Seq could detect pregnancy-related plasma cfRNA. **a**, The expression of placenta-specific mRNA or lncRNA in plasma during pregnancy. **b**, The expression of placenta-specific miRNA (C17MC cluster) in plasma during pregnancy. **c**, TPM of PALM-Seq and ΔCT of RT-qPCR normalized to *B2M* of *CGA, CSH1, CSH2, PLAC4* and *S100A8*. **d**, RPM of PALM-Seq and ΔCT of RT-qPCR normalized to RN7SL of mir-526b-5p, mir-519d-3p, mir-515-5p, mir-512-3p and mir-454-3p. N: nonpregnant females; T1: females at 12 gestational week; T2: females at 24 gestational week; T3: females at 36 gestational week. N=4.

**Figure 6.** Usage of PALM-Seq in amniotic fluid cfRNA and RNA from leukocyte or placenta. **a**, Average ratio (n=4) for each type of RNA in different samples. **b-d**, Principal components analysis for mRNA (**b**), lncRNA (**c**) and miRNA (**d**). **e**, 100 differently expressed miRNAs between leukocyte and placenta by PALM-Seq. **f**, 200 differently expressed mRNAs or lncRNAs between leukocyte and placenta by PALM-Seq.

## Discussion

Although the kits designed for total RNA such as Ribo-Zero Gold Kit NuGen’s^13^ or RNA-Seq Ovation System Kit^14^, were used before, they are not proper for cell-free RNA. Now, Small RNA Seq, SMARTer Seq and ScriptSeq are considered as optimized choices^15,22–24^. However, Small RNA Seq method requires 5’ phosphorylated and 3’ hydroxyl^36^, and pretreatment for low input of samples with Tobacco Acid Pyrophosphatase (TAP) or RNA 5’ Pyrophosphohydrolase (RppH) and T4 PNK may cause significant loss. SMARTer Seq and ScriptSeq could only capture large fragments, but small fragments are the main fraction of cfRNA, and they also may not be used for long-stored samples. There is no suitable method to capture small RNA and long RNA at the same time. Additionally, these kits are also expensive for large scale usage.

In contrast, PALM-Seq is a flexible, low cost method which could cover most of RNA in plasma except mRNA fragment with 5’ Cap. Compared to commercial RNA-Seq kit, many details can be easily modified to capture certain fractions without additional purification. Different terminal modifications might be distinguished by the usage of T4 PNK, TAP or RppH; however, if TAP or RppH is used, buffer should be optimized. Different target depletion could also be achieved just by change DNA oligo set. If certain length of fragments was required, size selection could be used after PCR amplification.

RNase H based target depletion^20^ is recruited here because it does not need certain kit, and there is no sequence restriction as in CRISPR-based method (PAM sequence NGG is necessary for Cas9) used by SMARTer Seq. RNase H based target depletion method was not limited by fragment length as well (DSN-lite might have problems when the fragments were too short). Removing rRNA by magnetic bead capture was also not practical for cfRNA due to severely lost. CATS-Seq^11^ (which shared the same principle of Clontech SMARTer smRNA-Seq Kit) which is similar to PALM-Seq, is not compatible for RNase H method, because the digestion production could not be removed and would be conjunct with 5’ adaptor through template-switch.

PALM-Seq is also time-saving and easy to handle. One could finish library preparation for 24 samples in 10 hours or for 32 samples in 12 hours without multichannel pipettors or purification workstations, and it might also be used in automatic production line. In a word PALM-Seq is a powerful sequencing method suitable for large scale cfRNA study or clinical application.

## Acknowledgements

We are grateful to volunteers who donated their samples to this study. We appreciate colleagues at BGI-Shenzhen for sequencing. This project is supported by the National Natural Science Foundation of China (No.81300075), the Natural Science Foundation of Guangdong Province (No. 2014A030313795), the Shenzhen Municipal Government of China (No.JCYJ20170412152854656, JCYJ20180703093402288), the Shenzhen Peacock Plan (No. KQTD20150330171505310).

## Author Contributions

W.J.W., F.C., and S.Z. conceived and designed the project. X.Y., S.Z., J.Z., Y.X., Z.L., H.C., J.S., L.L., C.G., collected samples and extracted plasma. X.Y. and Z.L. constructed PALM-Seq libraries and carried out other experiments. T.W., Y.X., Q.Z., and J.X. wrote the PALM-Seq analysis pipeline and analyzed PALM-Seq data. W.J.W., X.X., J.W., and H.Y. analyzed and interpreted all data. X.Y. and T.W. wrote the manuscript. W.J.W., F.C., and S.Z. revised the manuscript. All authors read and approved the manuscript for submission.

## Ethical compliance

We complied with all relevant ethical regulations.

## Code availability

The custom codes used to analyze PALM-Seq data are available from https://github.com/wonderful1/PALM-Seq-cfRNA

## Data availability

All the sequencing datasets generated in this study, including PALM-Seq test (with/without PNKT, with/ without TD), PALM-Seq for plasma from pregnant or nonpregnant women and PALM-Seq for amniotic fluid cfRNA or RNA from leukocytes and placentas have been deposited in CNSA (https://db.cngb.org/cnsa/) with accession CNP0000506.

